# CycleDesigner: Leveraging RFdiffusion and HighFold to Design Cyclic Peptide Binders for Specific Targets

**DOI:** 10.1101/2024.11.27.625581

**Authors:** Chenhao Zhang, Zhenyu Xu, Kang Lin, Chengyun Zhang, Wen Xu, Hongliang Duan

## Abstract

Cyclic peptides are potentially therapeutic in clinical applications, due to their great stability and activity. Yet, designing and identifying potential cyclic peptide binders targeting specific targets remains a formidable challenge, entailing significant time and resources. In this study, we modified the powerful RFdiffusion model to allow the cyclic peptide structure identification and integrated it with ProteinMPNN and HighFold to design binders for specific targets. This innovative approach, termed cycledesigner, was followed by a series of scoring functions that efficiently screen. With the combination of effective cyclic peptide design and screening, our study aims to further broaden the scope of cyclic peptide binder design.

## INTRODUCTION

Cyclic peptides represent a unique class of biomolecules characterized by their cyclic backbone structure. Compared to linear peptides, cyclic peptides exhibit notable advantages, such as enhanced stability against enzymatic degradation, stronger binding affinity, and higher target specificity[1-3]. These features make cyclic peptides promising candidates for therapeutic development, especially in targeting protein-protein interactions and other challenging molecular interfaces. However, the design of cyclic peptides is inherently complex due to their restricted conformational space and the intricate relationship between sequence and structure[4].

Traditional cyclic peptide design approaches often rely on extensive experimental filtering or heuristic computational methods, which can be both time-intensive and resource-demanding[5-7]. Recent advancements in computational modeling, particularly deep learning, have paved the way for novel strategies in de novo peptide design. RFdiffusion[8], a generative model based on probabilistic diffusion, has demonstrated remarkable success in designing proteins and protein scaffolds by exploring sequence-structure relationships in high-dimensional space. However, applying this approach to cyclic peptides remains a significant challenge. Limited availability of cyclic peptide data constrains large-scale training[9], and existing models often adapt pre-trained frameworks by modifying input features to accommodate specific cyclic peptide tasks[10].

In this study, we extend RFdiffusion to address the unique challenges of cyclic peptide design. By modifying the model’s input features to reflect the distinct structural characteristics of cyclic peptides, our approach enables the generation of novel cyclic peptide backbones and sequences for specific targets, without requiring extensive retraining on cyclic peptide datasets. We called it CycleDesigner. Through a series of computational experiments, we evaluate the performance of the CycleDesigner model and demonstrate its ability to generate structurally stable cyclic peptides. This work underscores the potential of advanced diffusion models and other deep learning techniques to revolutionize cyclic peptide design, bridging the gap between computational predictions and experimental validation.

## METHODS AND MATERIALS

### Cyclic peptide complex dataset

The data used in this study were sourced from the Protein Data Bank (PDB)[11], a publicly accessible repository of three-dimensional structural data for biomolecules. Specifically, we utilized the dataset of HighFold[12], focusing exclusively on single-chain protein structures. This restriction aligns with the requirements of RFdiffusion, which supports only single-chain protein inputs. As a result, multichain assemblies, including dimers, trimers and higher-order oligomers, were excluded from the dataset. For target structures with issues such as missing amino acids, we employed PDBfixer[13] to repair the proteins. This ensures the integrity of the single-chain protein structures and their compatibility with the input constraints of RFdiffusion. This preprocessing step was critical for maintaining the continuity and accuracy of the input protein chains, thereby enabling effective cyclic peptide binder design.

### Model

#### Computational Environment and Framework

We use Docker containers to establish an isolated and compatible computational environment[14]. Docker enabled seamless execution of RFdiffusion by encapsulating all dependencies within a standardized container, ensuring consistency across different computational setups. Docker also makes full use of GPU cuda acceleration and high-performance computing to achieve efficient modeling of cyclic peptides. This approach provided a robust and flexible framework for executing RFdiffusion without modifying the local server configuration, streamlining the computational process for cyclic peptide design.

#### Input Data Preprocessing

Before experiments, a series of data preprocessing steps were applied to ensure compatibility of the PDB files with the modified workflow. Firstly, ligands and other non-protein entities were removed from the PDB files, leaving only the target protein structure. This step simplified the input data, focusing solely on the protein chain required for RFdiffusion. Then RFdiffusion gets information about chain length, sequence identifiers, and residue indices derived from the raw PDB files. This information served as the basis for downstream modeling and sequence-structure alignment. We also retained relevant metadata of cyclic peptide complexes, such as ligand lengths and computed distances between ligands and the target protein binding site. This information was incorporated into the model as input features for constructing the positional encoding matrix, which is critical for guiding the cyclic peptide generation process.

#### Modification of Positional Encoding for Cyclic Peptides

The pre-trained RFdiffusion model incorporates positional encoding in its input features to represent spatial relationships among amino acid residues in linear protein chains. However, this linear encoding is unsuitable for cyclic peptides due to their circular topology. To address this limitation, we implemented the following modifications. We constructed a matrix of Cyclic relative position, using the input residue index (idx) from the linear representation and reflecting the circular topology of cyclic peptides. This matrix encodes the relative distances between each pair of amino acids, with the values representing the number of intervening residues. Additionally, the signs indicate the relative direction within the circular structure (clockwise or counterclockwise). The resulting matrix has a size of N×N, where N is the number of residues in the cyclic peptide. We then replaced the constructed cyclic relative position matrix replaced the default linear positional encoding in the input feature set of RFdiffusion. This modification allowed the model to effectively interpret and manipulate the cyclic topology without requiring additional retraining of the pre-trained framework.

#### Structure Output and Downstream Processing

The output of RFdiffusion is a cyclic peptide backbone in PDB format, accurately representing the cyclic topology and maintaining realistic spatial constraints. The downstream processing integrates the following steps: sequence generation with ProteinMPNN[15] and structure refinement with HighFold[12].

We employed the structure-based graph neural network model ProteinMPNN for sequence generation. ProteinMPNN specializes in generating amino acid sequences based on the 3D structural features of proteins, including N, Cα, C, O, and virtual Cβ atoms. The encoder comprises multiple layers with hidden dimensions of 128. For each residue iii, its node features are updated by integrating features from neighboring residues and edge connections through a message-passing mechanism. This iterative process ensures that structural and relational information is effectively captured. The decoder translates the encoded structural information into residue probabilities for each position. For tasks involving fixed-target ligand generation, ProteinMPNN incorporates known sequence segments as background constraints. These are combined with the structural features of the unknown regions to infer the sequence. The model is trained by minimizing the cross-entropy loss for each residue, quantifying the deviation between predicted and original sequences.

The newly generated cyclic peptide sequence is fed into HighFold, a deep learning model based on AlphaFold[16, 17] for predicting the 3D structures of cyclic peptides and their complexes to further explore the plausibility of designed backbones and sequences. HighFold introduces the Cyclic Position Offset Encoding Matrix (CycPOEM) to adapt AlphaFold’s positional encoding for cyclic peptide topology. This enhancement ensures accurate prediction of the cyclic peptide structure and its interaction with target proteins. By combining RFdiffusion, ProteinMPNN, and HighFold, our workflow efficiently generates novel cyclic peptide binders for specific targets and predicts their sequences and structures, offering a powerful pipeline for cyclic peptide binder design and analysis.

In this study, we use Rosetta’s energy analyzer[18] to evaluate the stability and binding affinity of the generated cyclic peptide-protein complexes. To filter and select high-confidence designs, we apply the energy metric dGseparated/dSASA×100. This value serves as an indicator of the peptide’s binding efficiency, helping us identify structures with favorable energy characteristics for further analysis and experimental validation.

#### Hotspot Identification

In the context of ligand-target interactions, hotspots are key residues at the binding site that play a critical role in molecular recognition and binding affinity[18-20].In this study, we defined hotspots as target residues within a 5 Å threshold of the ligand. Two strategies were considered for computing the shortest distance between ligand atoms and target residues: one-to-one and one-to-many (Adopted) Strategy. For the approach of one-to-one, for each ligand residue, all distances to target residues are computed, and the residue with the shortest distance is selected as a hotspot. This approach often results in a limited or empty hotspot set, particularly for ligands with sparse or irregular interaction patterns. Insufficient hotspot guidance can lead to off-target backbone generation during modeling, where the backbone deviates from the intended binding site due to the lack of spatial constraints. For the one-to-many approach, distances from each ligand residue to all target residues are computed. Any target residue within the 5 Å threshold is considered a hotspot. Reducing the risk of insufficient hotspot identification by ensuring a more comprehensive selection of residues within the interaction interface. Capturing a broader representation of the binding interface offers the model richer spatial constraints. Aligns generated backbones more closely with the binding site, increasing the reliability and applicability of the model outputs for downstream tasks.

### Experiment settings

All the experiments run on an Ubuntu (version 20.04.2) workstation with hardware configurations as follows: Intel i9 CPU (16 cores), RTX 3090 GPU*2 (24GB*2 Video.

RAM in total), 64GB Memory and 6TB Hard Disk. RFdiffusion is developed and implemented based on docker containers.

## RESULTS AND DISCUSSION

### Effect of Diffusion Steps (T) on Cyclic Peptide Design

The parameter diffuser.T, representing the number of diffusion steps, significantly influences the accuracy of the CycleDesigner outputs. While increasing diffusion steps generally enhances output accuracy, our experiments demonstrated a distinct trend for cyclic peptide backbone generation. We evaluated different diffusion steps by redesigning all targets in the dataset, selecting the top-1 backbone generated by CycleDesigner, generating the top-1 sequence with ProteinMPNN, and predicting the 3D structure with HighFold. Key metrics included pLDDT for local structural confidence, RMSD for alignment accuracy, and ipTM for assessing the quality of predicted complexes. The results showed that the pLDDT metric remained relatively consistent across diffusion steps, with T=25 and T=30 yielding better performance. Specifically, pLDDT values at T=25 were more enriched at higher levels. RMSD values showed an average of 2.151 Å at the default T=20 and improved with increasing steps, but performance degraded at T≥30 with an average RMSD of 2.212 Å. The scatterplot analysis revealed that RMSD values at T=25 were more concentrated and enriched at lower levels. Similarly, ipTM values peaked at T=25 with an average of 0.541, compared to 0.527 at T=20, and decreased to 0.511 at T=35. Notably, CycleDesigner with T=25 also produced backbones with the highest number of ipTM results above 0.8, indicating a higher likelihood of generating structures with binding activity. These results highlight a unique behavior for cyclic peptides, where increasing diffusion steps does not always improve accuracy. This is likely due to the model’s bias toward generating α-helical secondary structures, which are less prominent in cyclic peptides, where loop structures dominate. Higher diffusion steps exacerbate this bias, reducing structural accuracy for cyclic peptides.

Therefore, diffuser.T=25 was selected as the optimal diffusion step for cyclic peptide design in our workflow, striking a balance between model performance and structural relevance.

### Cyclic peptide binders designed using Rfdiffusion for specific targets

We performed the design of cyclic peptide ligands for 23 targets. Using CycleDesigner, we generated five cyclic peptide backbones for each target, resulting in a total of 115 backbones. These backbones were then processed through ProteinMPNN, with each backbone producing five novel sequences, yielding 575 new cyclic peptide sequences. The structural prediction of all 575 sequence designs was conducted using HighFold, generating 2,875 unique 3D structures of cyclic peptide-target complexes.

Among these complexes, the average pLDDT reached 91.263, while pTM, ipTM, and iPAE scores were 0.912, 0.533, and 0.465, respectively. Cyclic peptides with lengths of 7 and 14 residues were most prevalent, accounting for 625 and 1,500 structures, respectively. These results highlight the capability of the workflow to generate a large number of high-confidence cyclic peptide designs tailored for specific targets.

### The screening for potential cyclic peptide binders

We filtered the 2,875 cyclic peptide-target complex structures to identify candidates for subsequent energy function calculations. The filtering criteria included iPAE<0.35, ipTM>0.5, pLDDT>80, and RMSD to input backbone <3.5. This process yielded 305 high-quality designs.

After filtering, the average pLDDT of the selected structures significantly improved, exceeding 95, with most results surpassing 90. The ipTM, a critical indicator of complex quality, showed an even greater enhancement, with an average increase of 20%, reaching nearly 0.8. However, ipTM values were distributed between 0.7 and 0.8 before screening. The iPAE also improved markedly, with an average dropping below 0.3. The RMSD to the backbone, which already performed well, showed slight improvement. Its distribution exhibited two dense regions, one below 1 Å and another between 2 and 3 Å. This bimodal distribution may result from the variability in cyclic peptide lengths, as longer sequences tend to present greater challenges in achieving low RMSD after sequence design.

These findings underscore the effectiveness of the filtering process in finding high-quality cyclic peptide designs for further analysis.We performed energy function calculations on the filtered structures using Rosetta’s energy_analyze tool. Based on the criterion dGseparated/dSASA×100, 245 structures were selected. Figure 5A illustrates the best-performing designs for each target meeting this criterion. The RMSD distribution of all selected structures shows two prominent clusters around 1 Å and 2.5 Å. Among these, 74 structures exhibited dGseparated/dSASA×100<−1.5. Interestingly, there was no linear correlation between ipTM and dGseparated/dSASA×100. However, as ipTM increased, the enrichment region of dGseparated/dSASA×100 shifted slightly downward, suggesting a modest degree of correlation (Figure 5E). These results highlight the correlation between binding affinity metrics and structural quality indicators, offering valuable insights for further refinement and evaluation of cyclic peptide designs.

**Figure 1.**
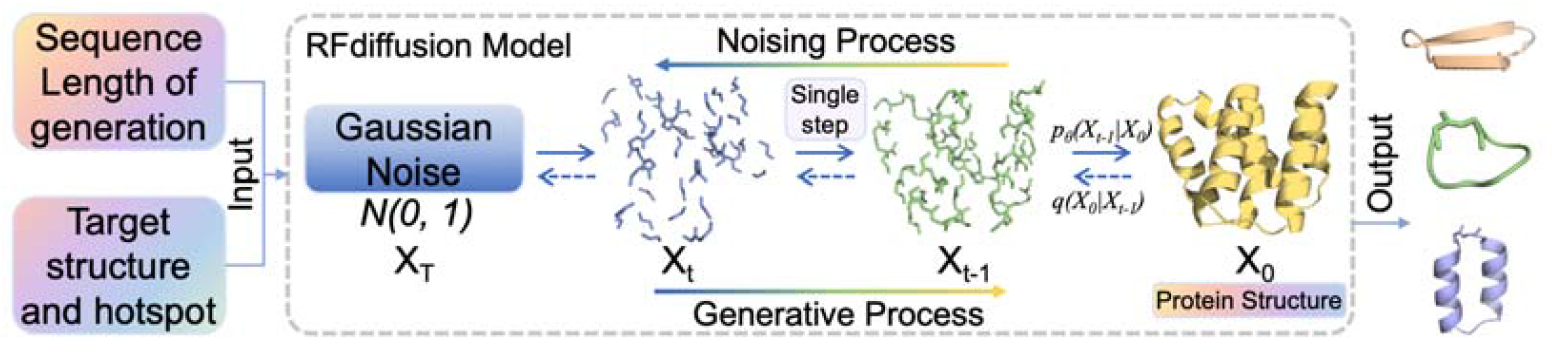
The illustration of modified RFdiffusion. The model requires the target protein structure file and designated hotspot regions as inputs. Additionally, the desired length of the cyclic peptide sequence must be specified.

**Figure 2.**
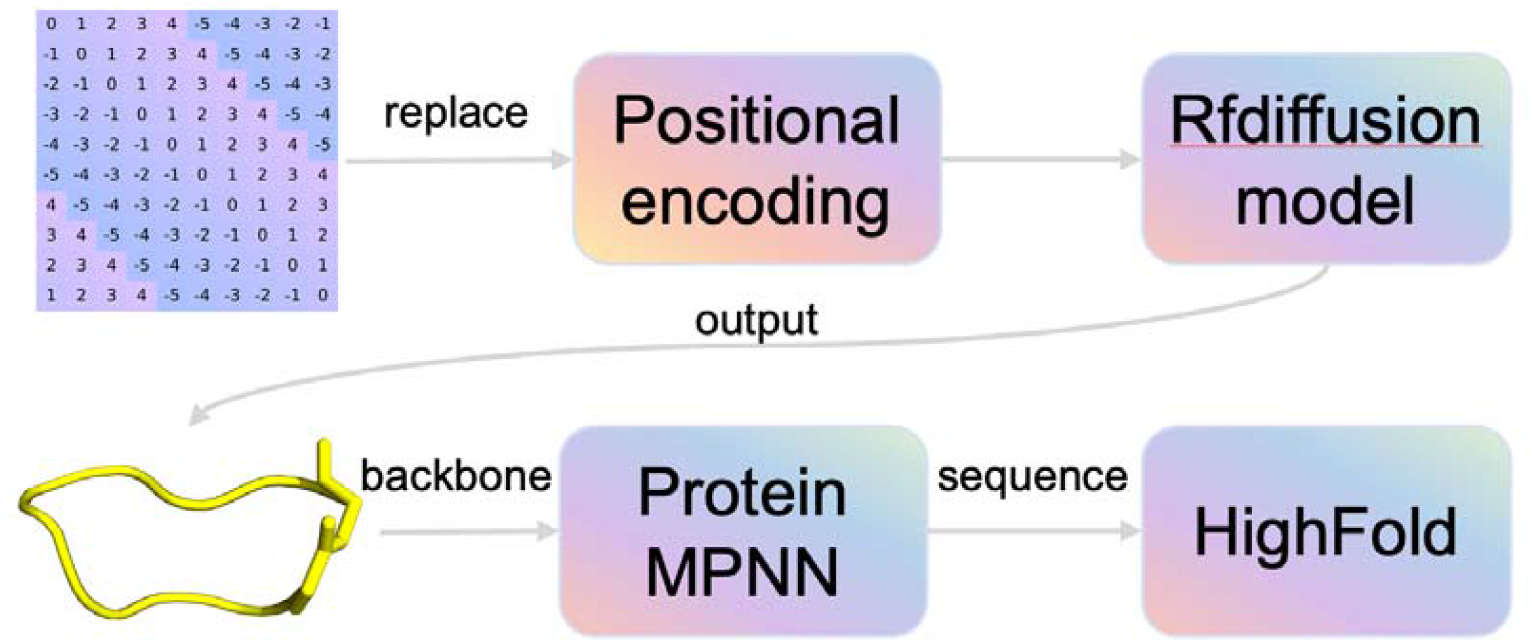
The complete workflow for designing cyclic peptides using RFdiffusion. By replacing the default relative position coding, RFdiffusion was able to generate a cyclic peptide backbone for specific targets. The backbone file was input into ProteinMPNN to obtain a new sequence. Finally, HighFold was used to predict the designed cyclic peptide structure.

**Figure 3.**
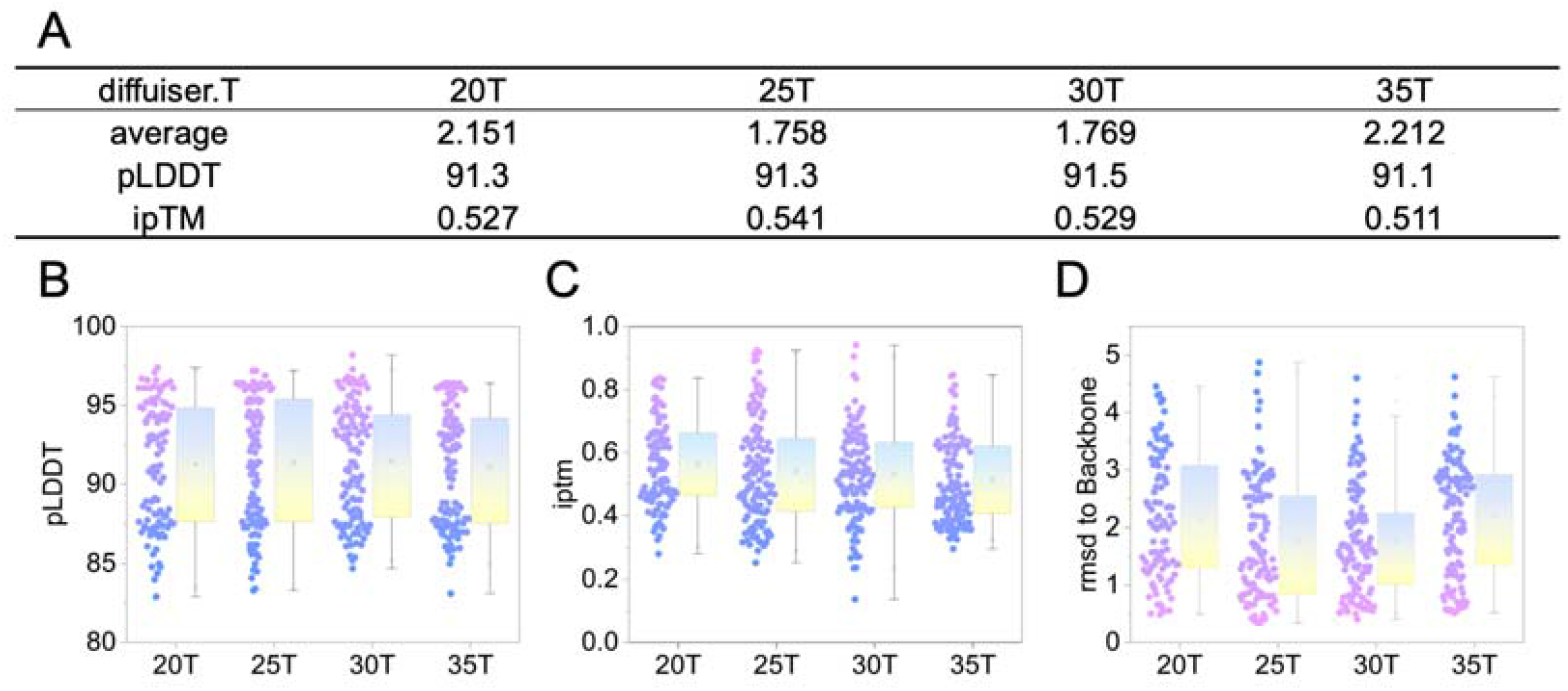
The results of different diffusion steps on cyclic peptide design. **a**. Comparison of important indicators of complexes with different diffusion steps. **b**. pLDDT distribution of different diffusion steps. **c**. ipTM distribution of different diffusion steps. **d**. RMSD to backbone distribution of different diffusion steps.

**Figure 4.**
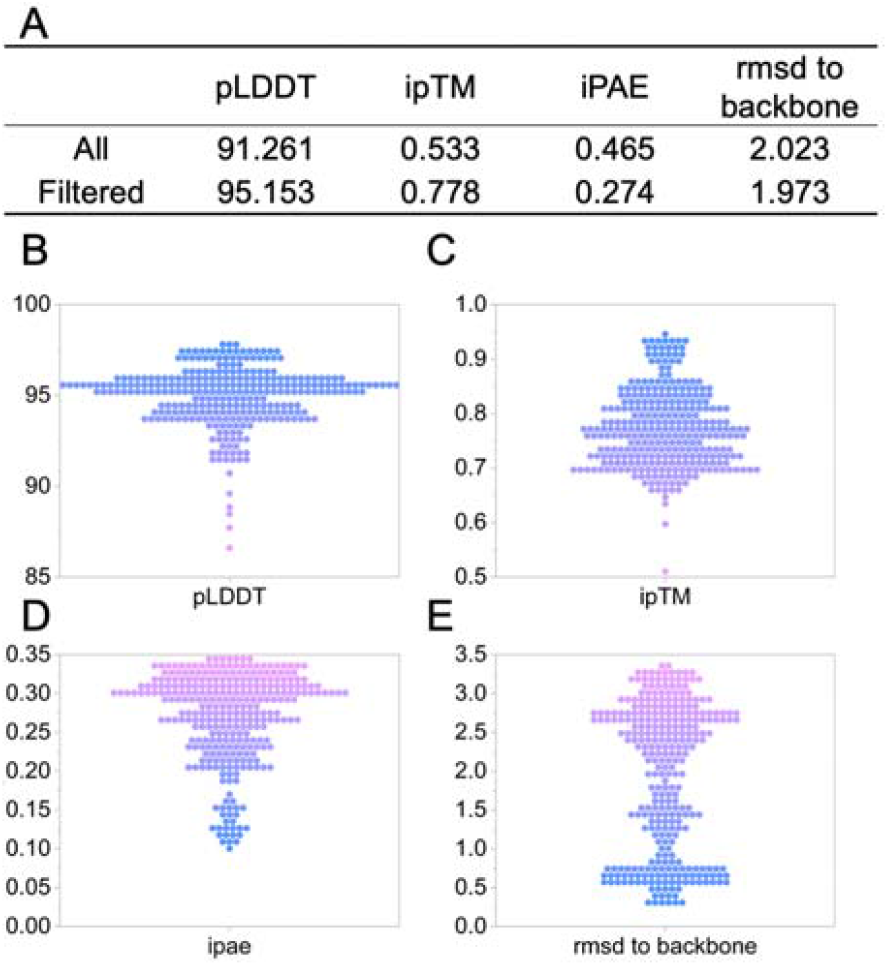
Preliminary filtering results of alphafold related indicators. **a**. Comparison of results before and after filtering. **b**. pLDDT distribution after filtering. **c**. ipTM distribution after filtering. **d**. iPAE distribution after filtering. **E**. RMSD to backbone distribution after filtering.

**Figure 5.**
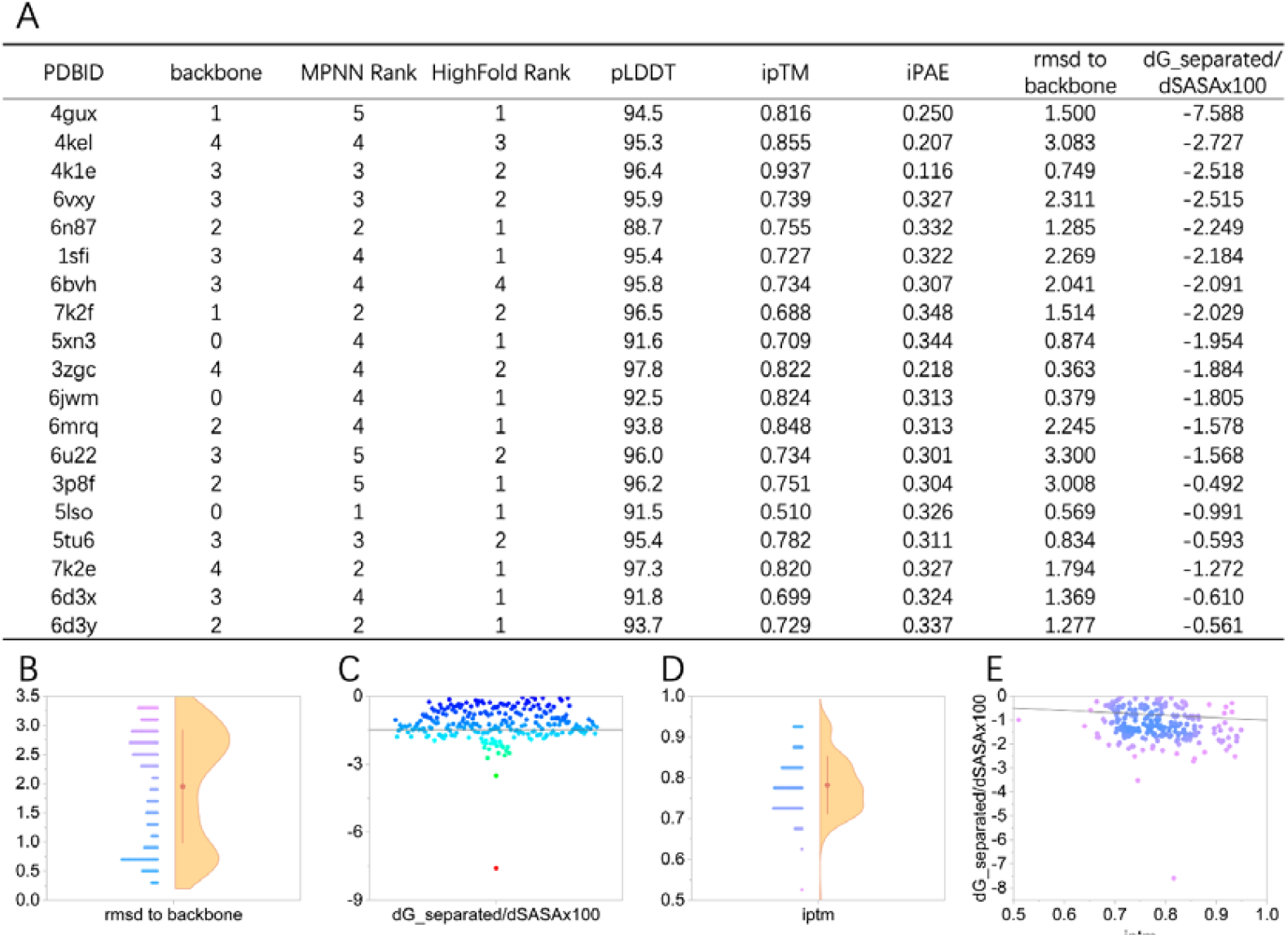
The filtering results using Rosetta energy analysis. **a**. Top 1 results of all targets that meet the filtering criteria. **b**. RMSD to backbone distribution of results after screening. **c**. dGseparated/dSASA×100 distribution of results after screening. **d**. ipTM distribution of results after screening. **e**. The correlation between dGseparated/dSASA×100 and ipTM.

Among the top 1 structures for all selected targets, we observed that the designed cyclic peptide ligands exhibited spatial positions similar to their natural counterparts. However, their 3D structures and sequences were significantly different. In cases such as 4gux and 6mrq, which involve longer cyclic peptide sequences, the structural similarity was particularly pronounced in the contact regions. Conversely, other regions displayed notable differences, likely due to the generative tendencies of the model (Figures 6 and 7).

**Figure 6.**
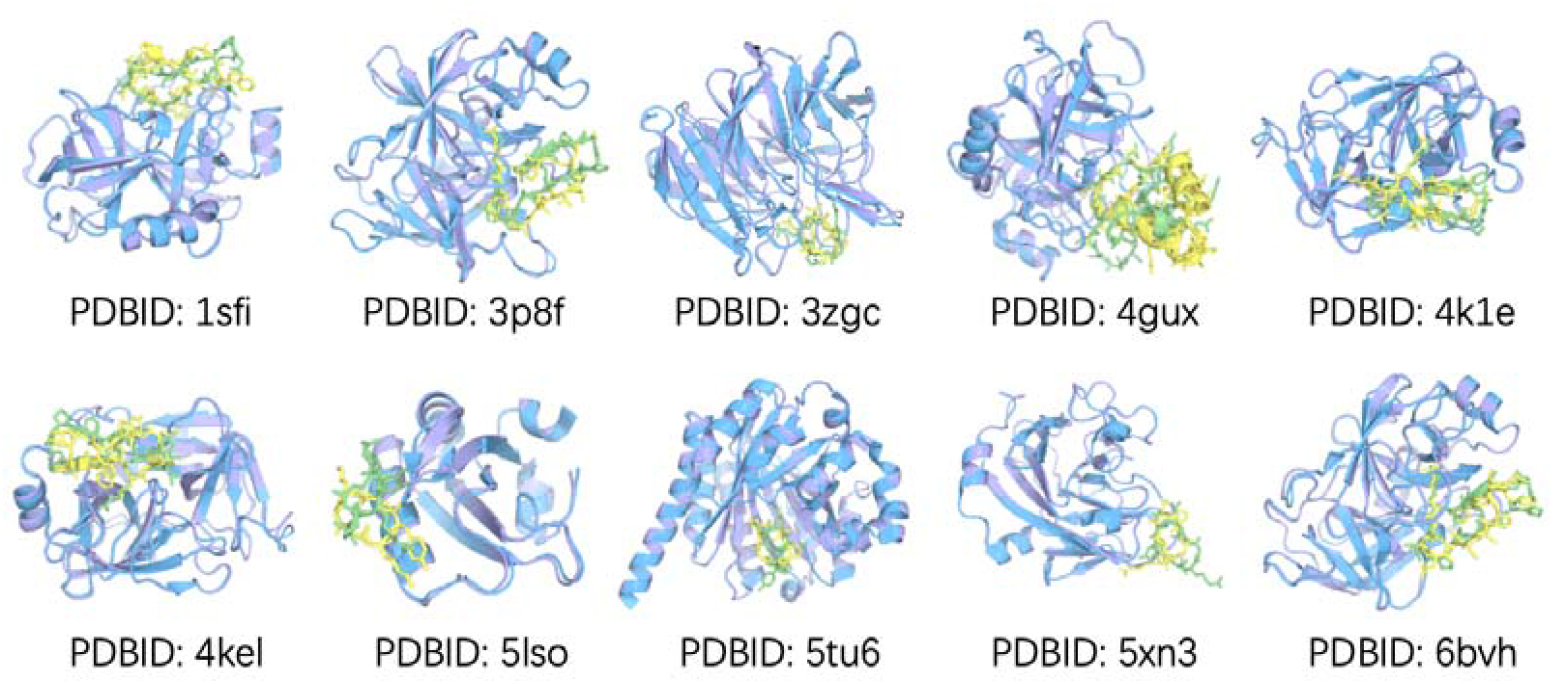
The comparisons of top 1 designed structures and native structures.

**Figure 7.**
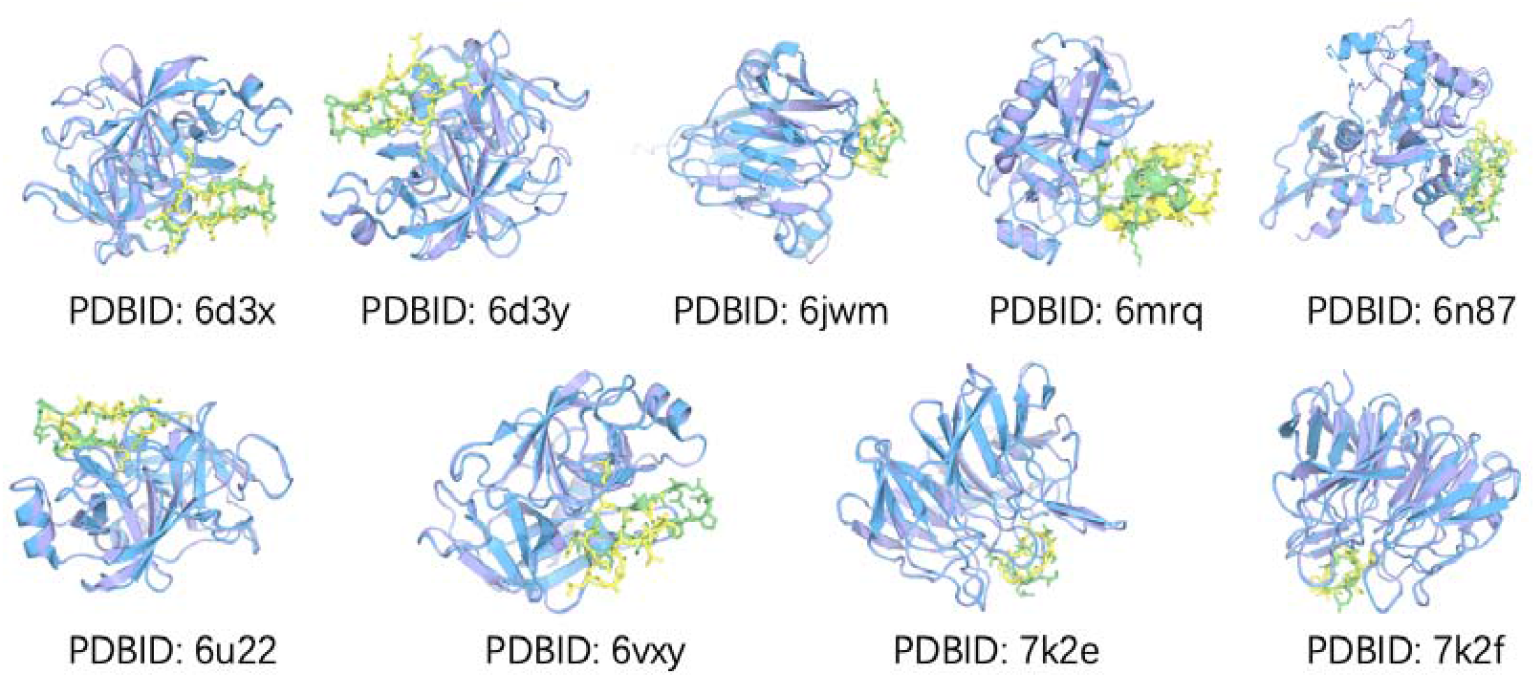
The comparisons of top 1 designed structures and native structures.

**Figure 7.**
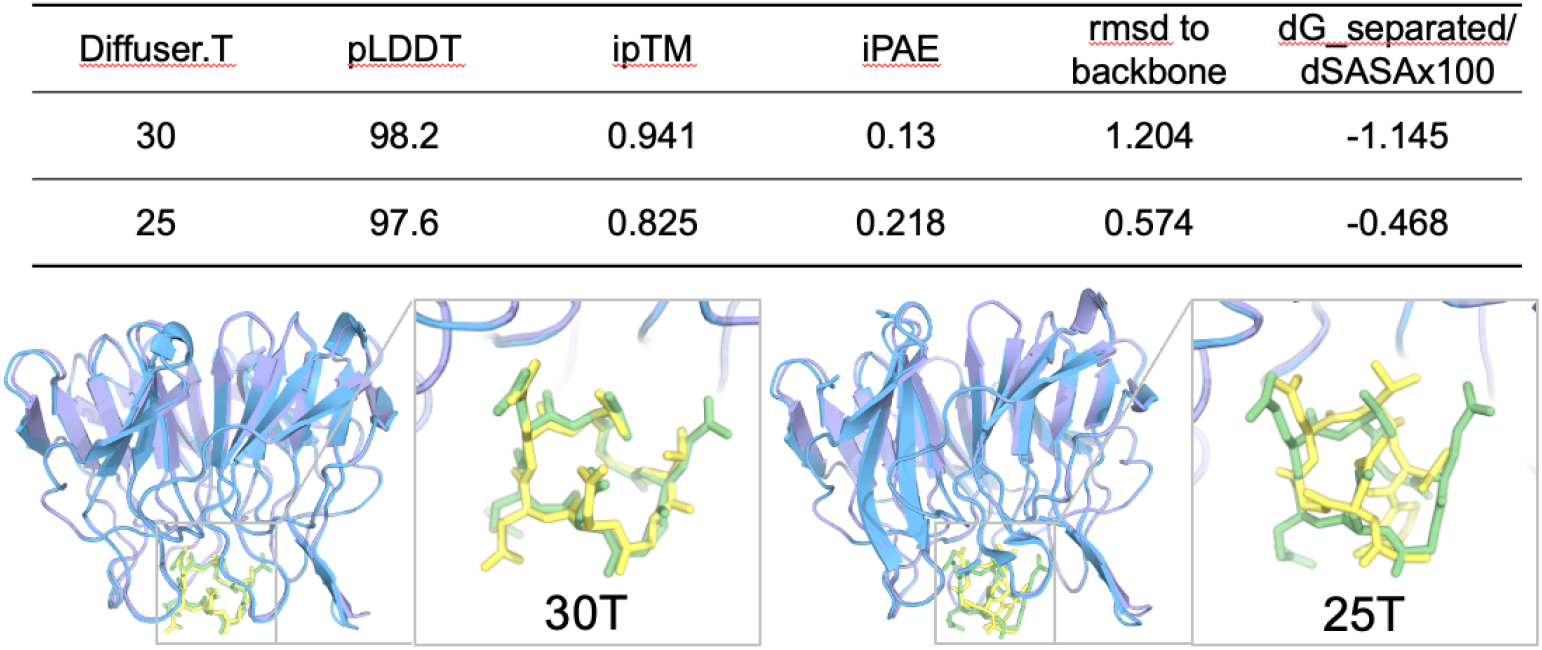
Comparison of various indicators of target 3zgc under different parameters, and comparison between the designed structures and natural structures under different parameters.

These findings highlight the model’s ability to retain spatial alignment with native ligands while introducing diversity in sequence and non-contacting structural elements, potentially broadening the functional scope of the designed cyclic peptides.

### Parameter adjustment for diffrenret targets in cyclic peptide design

In exploring the impact of diffusion steps, we found that different targets may have distinct optimal parameters. For example, with 3ZGC, the best results were achieved at 30 diffusion steps. At this setting, three identical amino acid sequences were generated, showing high spatial overlap. Other metrics also performed exceptionally, with pLDDT reaching 98.2, ipTM 0.941, RMSD to backbone 1.204, and dGseparated/dSASA×100 at -1.145. In contrast, at 25 diffusion steps-the parameter used in our batch processing—the best filtered results showed slightly lower metrics, including a pLDDT of 97.6, ipTM 0.825, iPAE 0.218, RMSD to backbone 0.573, and a Rosetta energy score of -0.468. Structurally, the design at 30 diffusion steps was also more similar to the native structure (Figure 8).

These findings indicate that cyclic peptide design results exhibit a degree of randomness. Selecting universally applicable parameters, such as 25 diffusion steps, increases the likelihood of obtaining good designs but may not always yield the optimal structures for specific targets. For certain targets, adjusting parameters can lead to designs with higher potential, underscoring the importance of flexibility in parameter selection for cyclic peptide design.

## CONCLUSION

This study demonstrates the potential of CycleDesigner for cyclic peptide design. By introducing cyclic positional encoding, we effectively overcome the limitations of traditional generative models based on linear encoding. This approach allows us to design high-accuracy cyclic peptide topologies without the need for retraining or fine-tuning. Through our comprehensive workflow, we generated over 2,800 novel cyclic peptide-protein complexes. Of these, 305 candiated were retained after passing rigorous structural and energy-based screening criteria, and finally 245 high-confidence cyclic peptide complexes met multiple standards for further exploration.

In addtion, while using universal parameters can streamline the process, the stochastic nature of cyclic peptide design suggests that adjusting parameters for specific targets can significantly improve outcomes. This balance between generality and specificity represents the next key challenge in cyclic peptide computational design.

Our work advances the computational toolkit for cyclic peptide design, enabling the efficient generation of novel cyclic peptide binders with favorable structural and energy properties, even in the absence of detailed ligand information. Future research should focus on experimental validation of the best candidates and further refinement of the models to enhance their applicability in a broad range of peptide-protein interaction environments.

## CODE AVAILABILITY

The code and scripts for running the model will be released at GitHub after publication.

### Author Biographies

**Chenhao Zhang** is a postgraduate student in the Artificial Intelligence Assisted Drug Discovery Institute and the Department of Pharmacy, Zhejiang University of Technology. His research interests are in bioinformatics, peptides discovery, machine learning and deep learning.

**Zhenyu Xu** is a researcher in the Shenzhen Highslab Therapeutics. Inc. His research interests are in bioinformatics, computational biology, machine learning and deep learning.

**Kang Lin** is a postgraduate student in the Artificial Intelligence Assisted Drug Discovery Institute and the Department of Pharmacy, Zhejiang University of Technology. His research interests are in bioinformatics, peptides discovery, machine learning and deep learning.

**Xu Wen** is a postgraduate student in the Artificial Intelligence Assisted Drug Discovery Institute and the Department of Pharmacy, Zhejiang University of Technology. Her research interests are in pharmacological data analysis, computational biology, machine learning and deep learning.

**Chengyun Zhang** is a researcher in the Artificial Intelligence Assisted Drug Discovery Institute and the Department of Pharmacy, Zhejiang University of Technology. Her research interests are in bioinformatics, computational biology, machine learning and deep learning.

**Hongliang Duan** is a professor in the Artificial Intelligence Assisted Drug Discovery Institute and the Department of Pharmacy, Zhejiang University of Technology. His research interests are in AI-driven drug discovery, machine learning and deep learning.

